# A Conserved Mechanism of APOBEC3 Relocalization by Herpesviral Ribonucleotide Reductase Large Subunits

**DOI:** 10.1101/765735

**Authors:** Adam Z. Cheng, Sofia Nóbrega de Moraes, Claire Attarian, Jaime Yockteng-Melgar, Matthew C. Jarvis, Matteo Biolatti, Ganna Galitska, Valentina Dell’Oste, Lori Frappier, Craig J. Bierle, Stephen A. Rice, Reuben S. Harris

## Abstract

An integral part of the antiviral innate immune response is the APOBEC3 family of single-stranded DNA cytosine deaminases, which inhibits virus replication through deamination-dependent and -independent activities. Viruses have evolved mechanisms to counteract these enzymes such as HIV-1 Vif-mediated formation of a ubiquitin ligase to degrade virus-restrictive APOBEC3 enzymes. A new example is Epstein-Barr virus (EBV) ribonucleotide reductase (RNR)-mediated inhibition of cellular APOBEC3B (A3B). The large subunit of the viral RNR, BORF2, causes A3B relocalization from the nucleus to cytoplasmic bodies and thereby protects viral DNA during lytic replication. Here, we use co-immunoprecipitation and immunofluorescent microscopy approaches to ask whether this mechanism is shared with the closely related γ-herpesvirus Kaposi’s sarcoma-associated herpesvirus (KSHV) and the more distantly related α-herpesvirus, herpes simplex virus-1 (HSV-1). The large RNR subunit of KSHV, ORF61, co-precipitated multiple APOBEC3s including A3B and APOBEC3A (A3A). KSHV ORF61 also caused relocalization of these two enzymes to perinuclear bodies (A3B) and to oblong cytoplasmic structures (A3A). The large RNR subunit of HSV-1, ICP6, also co-precipitated A3B and A3A and was sufficient to promote the relocalization of these enzymes from nuclear to cytoplasmic compartments. HSV-1 infection caused similar relocalization phenotypes that required ICP6. However, unlike the infectivity defects previously reported for BORF2-null EBV, ICP6 mutant HSV-1 showed normal growth rates and plaque phenotypes. These results combine to indicate that both γ- and α-herpesviruses use a conserved RNR-dependent mechanism to relocalize A3B and A3A and, further, suggest that HSV-1 possesses at least one additional mechanism to neutralize these antiviral enzymes.

**Importance:** The APOBEC3 family of DNA cytosine deaminases constitutes a vital innate immune defense against a range of different viruses. A novel counter-restriction mechanism has recently been uncovered for the γ-herpesvirus EBV, in which a subunit of the viral protein known to produce DNA building blocks (ribonucleotide reductase) causes A3B to relocalize from the nucleus to the cytosol. Here, we extend these observations with A3B to include a closely related γ-herpesvirus, KSHV, and to a more distantly related α-herpesvirus, HSV-1. These different viral ribonucleotide reductases also caused relocalization of A3A, which is 92% identical to A3B. These studies are important because they suggest a conserved mechanism of APOBEC3 evasion by large double-stranded DNA herpesviruses. Strategies to block this host-pathogen interaction may be effective for treating infections caused by these herpesviruses.

## Introduction

An important arm of the innate immune response lies in the APOBEC family of single-stranded DNA cytosine deaminases (1-3). Each of the seven human APOBEC3 (A3) enzymes, A3A-D and A3F-H, have been implicated in the restriction and mutation of a variety of different human viruses including retroviruses (HIV-1, HIV-2, HTLV-1) (4-8), endogenous retroviruses (HERV) (9, 10), hepadnaviruses (HBV) (11, 12), small DNA tumor viruses (HPV, JC/BK-PyV) (13-17), and most recently, the γ-herpesvirus Epstein-Barr Virus (EBV) (18, 19). It is difficult, if not impossible, to predict *a priori* which subset of APOBEC3 enzymes has the potential to engage a given virus and, furthermore, how that virus might counteract potentially restrictive A3 enzymes. For instance, the lentiviruses HIV-1 and HIV-2 encode an accessory protein called Vif that heterodimerizes with the cellular transcription co-factor CBF-β and recruits a cellular ubiquitin ligase complex to trigger the degradation of restrictive A3 enzymes (20, 21).

Human herpesviruses can be grouped into three distinct subfamilies (α, β, and γ; phylogeny shown in **Fig 1A**). Pathogenic α- and β-herpesviruses include herpes simplex virus type 1 (HSV-1) and cytomegalovirus (CMV), respectively, and the γ-herpesvirus subfamily includes EBV and Kaposi’s sarcoma-associated herpesvirus (KSHV). We recently identified an A3 counteraction mechanism for EBV (18). We demonstrated that the large subunit of the viral ribonucleotide reductase (RNR), BORF2, inhibits APOBEC3B (A3B) by directly binding and relocalizing it from the nucleus to the cytoplasmic compartment. This counteraction mechanism prevents the normally nuclear-localized A3B enzyme from deaminating viral genomic DNA cytosines to uracils during lytic replication. In the absence of BORF2, A3B inflicted C/G-to-T/A mutations in EBV genomes and reduced viral titers and infectivity. We also showed that the homologous protein from KSHV, ORF61, is similarly capable of binding and relocalizing A3B (18).

**Fig 1.**
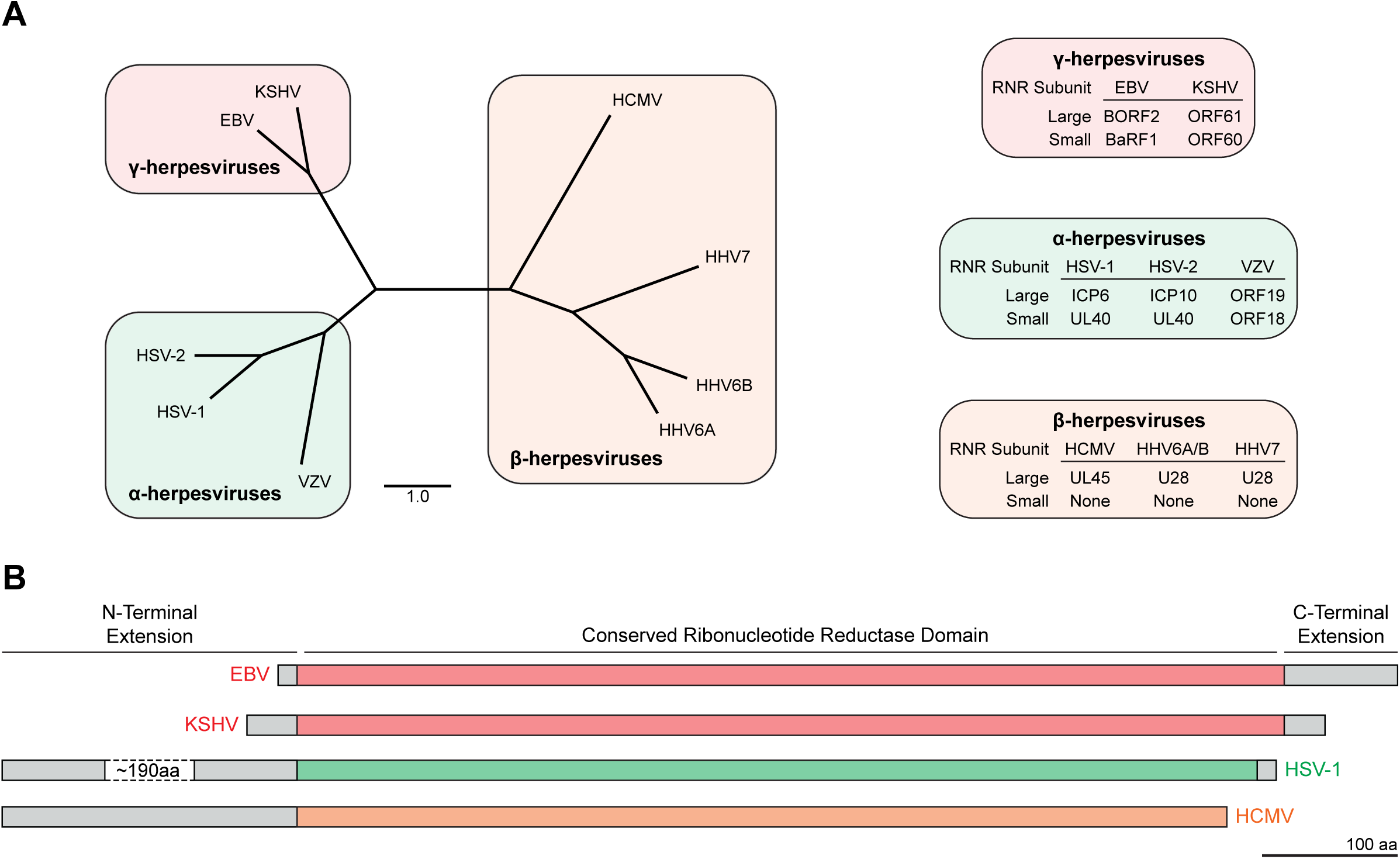
Herpesvirus ribonucleotide reductases conservation. **(A)** Amino acid sequences from ribonucleotide reductase large subunits were aligned using Multiple Sequence Comparison by Log-Expectation (MUSCLE) and phylogeny was constructed using neighbor-joining tree without distance corrections and scaled for equal branch lengths (scale bar = 1). Shaded boxes indicate herpesvirus subfamilies, which group closely to established phylogenetic trees. Protein names for human herpesvirus ribonucleotide reductase large and small subunits shown on the right. **(B)** Schematic of representative RNR large subunit polypeptides from α-, β-, and γ-herpesviruses with conserved core sequences (colored) and unique N- and C-terminal extensions (gray). Diagram is approximately to scale with a ∼190 amino acid portion of HSV-1 ICP6 omitted to fit the figure. Scale bar is 100 amino acids.

Here, we ask whether the viral RNR-mediated A3B counteraction mechanism is specific to γ-herpesviruses or more general-acting by assessing interactions between γ-herpesvirus BORF2/ORF61 and other human A3 enzymes and by determining whether the more distantly related α-herpesvirus HSV-1 has a similar A3 neutralization mechanism (RNR nomenclature in **Fig 1A** and protein domains depicted in **Fig 1B**). We found that, in addition to binding and relocalizing A3B, both BORF2 and ORF61 were also capable of co-immunoprecipitation and relocalization of A3A. Additionally, we found that the HSV-1 RNR large subunit ICP6 similarly binds and relocalizes both A3B and A3A. Overexpression studies showed that ICP6 alone is sufficient for A3B and A3A relocalization. Infection studies with wild-type and mutant viruses demonstrated that ICP6 mediates this relocalization activity in the context of infected cells and that no other viral protein is capable of this relocalization function. However, despite likely conservation of the A3B/A relocalization mechanism, the infectivity of ICP6 mutant HSV-1 was not affected by A3B or A3A suggesting the existence of a functionally redundant A3 neutralization mechanism.

## Results

### EBV BORF2 and KSHV ORF61 bind and relocalize both A3B and A3A

Our prior co-immunoprecipitation (co-IP) experiments indicated that EBV BORF2 interacts strongly with A3B and weakly with A3A and A3F (see Fig. 1c in Cheng *et al*. (18)). EBV BORF2 was both necessary and sufficient to relocalize A3B in a variety of different cell types including endogenous A3B in the AGS gastric carcinoma cell line and the M81 B cell line (18). However, our original studies did not address whether EBV BORF2 could functionally interact with and relocalize any of these related human A3 enzymes. We therefore performed immunofluorescent (IF) microscopy studies of U2OS cells overexpressing A3-mCherry constructs with either empty vector or BORF2-FLAG. As reported, A3B is nuclear, A3A has a cell-wide localization, A3H is cytoplasmic and nucleolar, and the other A3s are cytoplasmic (22-26). Also as expected, BORF2 caused a robust and complete relocalization of nuclear A3B to perinuclear aggregates (**Fig 2**). Interestingly, BORF2 co-expression with A3A led to the presence of novel linear elongated structures concomitant with normal A3A localization. The localization patterns of the other five A3s were unchanged by BORF2 co-expression. Small BORF2 punctate structures were also noted in all conditions including the mCherry control, which is likely due to transfected BORF2 interacting with endogenous A3B (previously shown to be elevated in U2OS (18)). Similar A3B and A3A relocalization patterns were evident in Vero cells except that A3A relocalization became whole-cell without elongated structures (**Supplementary Fig 1**).

**Fig 2.**
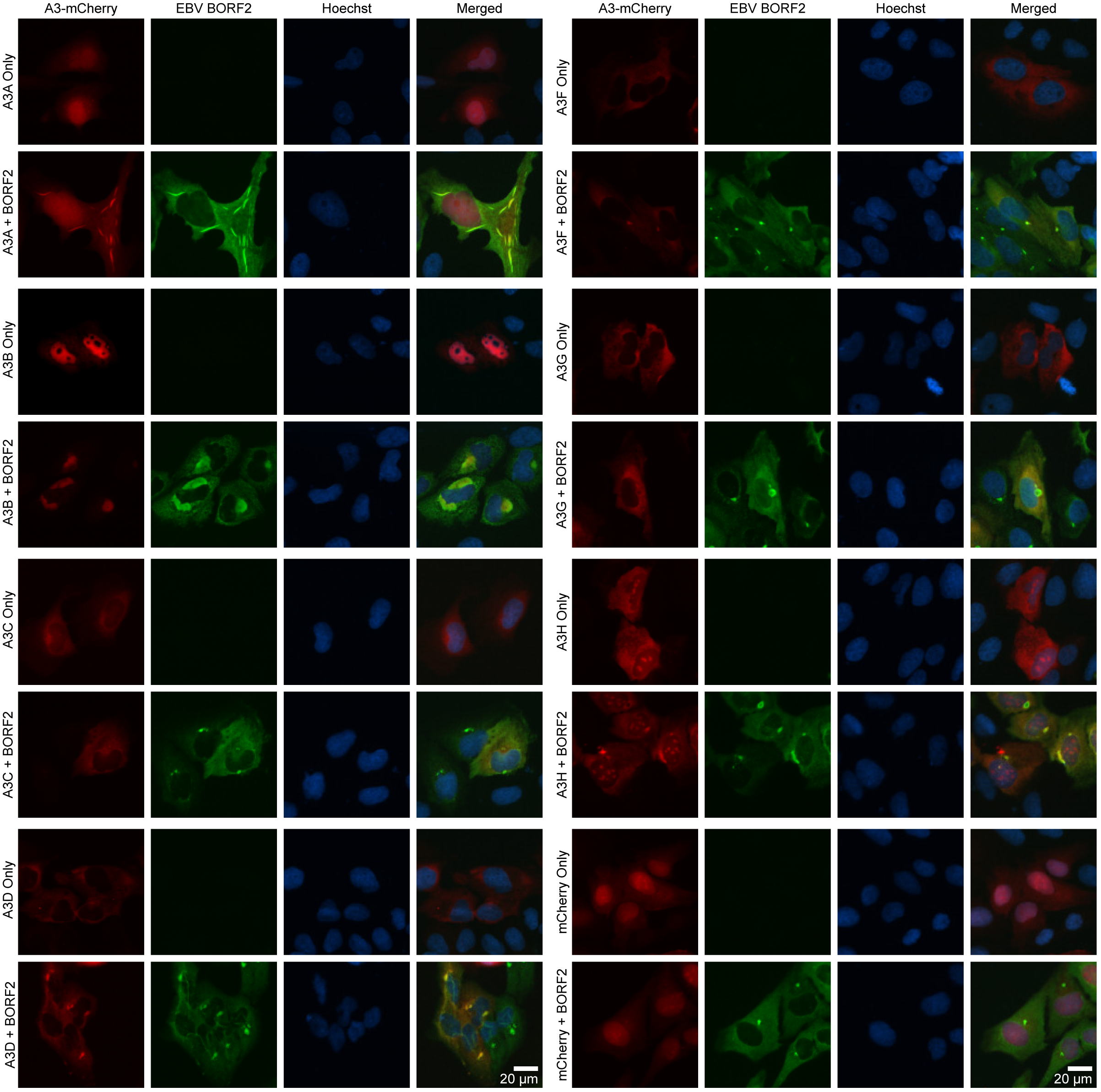
EBV BORF2 relocalizes A3B and A3A. Representative images of U2OS cells transfected with either A3-mCherry or BORF2-FLAG constructs. Cells were fixed 48 hours post-transfection, permeabilized, and stained with anti-FLAG antibody and Hoechst. A3 localization was compared in the presence and absence of EBV BORF2-FLAG co-transfection.

Like EBV BORF2, KSHV ORF61 was also shown to co-IP and relocalize A3B (18). However, our original studies did not examine the specificity of this interaction by comparing with related human A3 enzymes. We therefore used co-IP experiments to evaluate KSHV ORF61 interactions with a full panel of human A3 enzymes. ORF61-FLAG was co-expressed with A3-HA family members in 293T cells, subjected to anti-FLAG affinity purification, and analyzed by immunoblotting (**Fig 3A**). The ORF61-FLAG pulldown resulted in A3B recovery as described (18). In addition, the ORF61-FLAG IP also yielded a robust interaction with A3A and weaker interactions with A3D and A3F.

**Fig 3.**
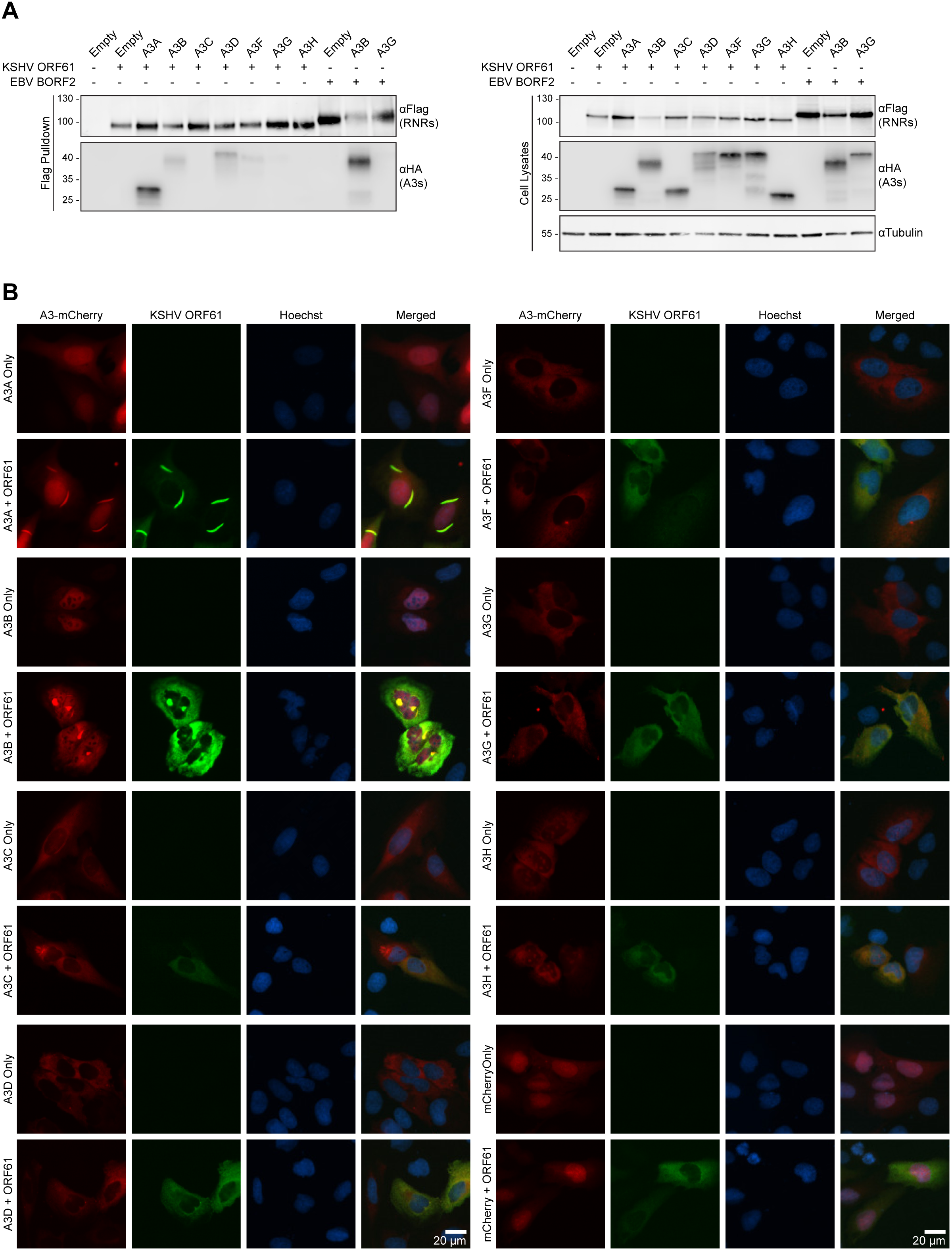
KSHV ORF61 relocalizes A3B and A3A. **(A)** Co-immunoprecipitation of transfected KSHV ORF61-FLAG with the indicated A3-HA constructs in 293T cells. Cells were lysed 48 hours post-transfection for anti-FLAG pulldown and resulting proteins were analyzed by immunoblot. EBV FLAG-BORF2 transfected with A3B and A3G were used as positive and negative co-IP controls, respectively. **(B)** Representative images of U2OS cells transfected with either A3-mCherry or FLAG-RNR constructs. Cells were fixed 48 hours post-transfection, permeabilized, and stained with anti-FLAG antibody and Hoechst. Co-transfection with A3B-mCherry and EBV BORF2-FLAG was used as positive controls for relocalization from nuclear to cytoplasmic aggregates. A3 localization was compared in the presence and absence of KSHV ORF61-FLAG co-transfection.

These KSHV ORF61-A3 interactions were then evaluated by IF microscopy experiments to look for changes in A3 localization in U2OS cells (**Fig 3B**). As expected (18), KSHV ORF61 caused A3B to relocalize to perinuclear bodies. Moreover, as above for BORF2 and A3A, ORF61 co-expression caused a portion of the cellular A3A to localize to intense elongated linear structures in the cytosolic compartment (**Fig 3B**). No other A3 proteins showed altered subcellular localization in these experiments. Similar IF microscopy observations were made using the same constructs in HeLa cells. These new results combined to indicate that both A3B and A3A are cellular targets of EBV BORF2 and KSHV ORF61. The potential relevance of these interactions to the pathogenesis of these viruses will be considered in the **Discussion**.

### HSV-1 ICP6 binds and relocalizes A3B and A3A

To test whether the RNR-mediated A3B/A relocalization mechanism is more broadly conserved, a series of co-IP experiments was done with the large RNR subunit of the pathogenic α-herpesvirus HSV-1, ICP6. FLAG-ICP6 was co-expressed with each of the seven different HA-tagged human A3s in 293T cells and subjected to anti-FLAG IP as above. The EBV BORF2-A3B interaction was used as a positive control and BORF2-A3G as a negative control to be able to compare the relative strengths of pulldowns between RNRs and A3s. HSV-1 ICP6 showed a strong interaction with A3A and weaker, but detectable, interactions with A3B, A3C, and A3D (**Fig 4A**).

**Fig 4.**
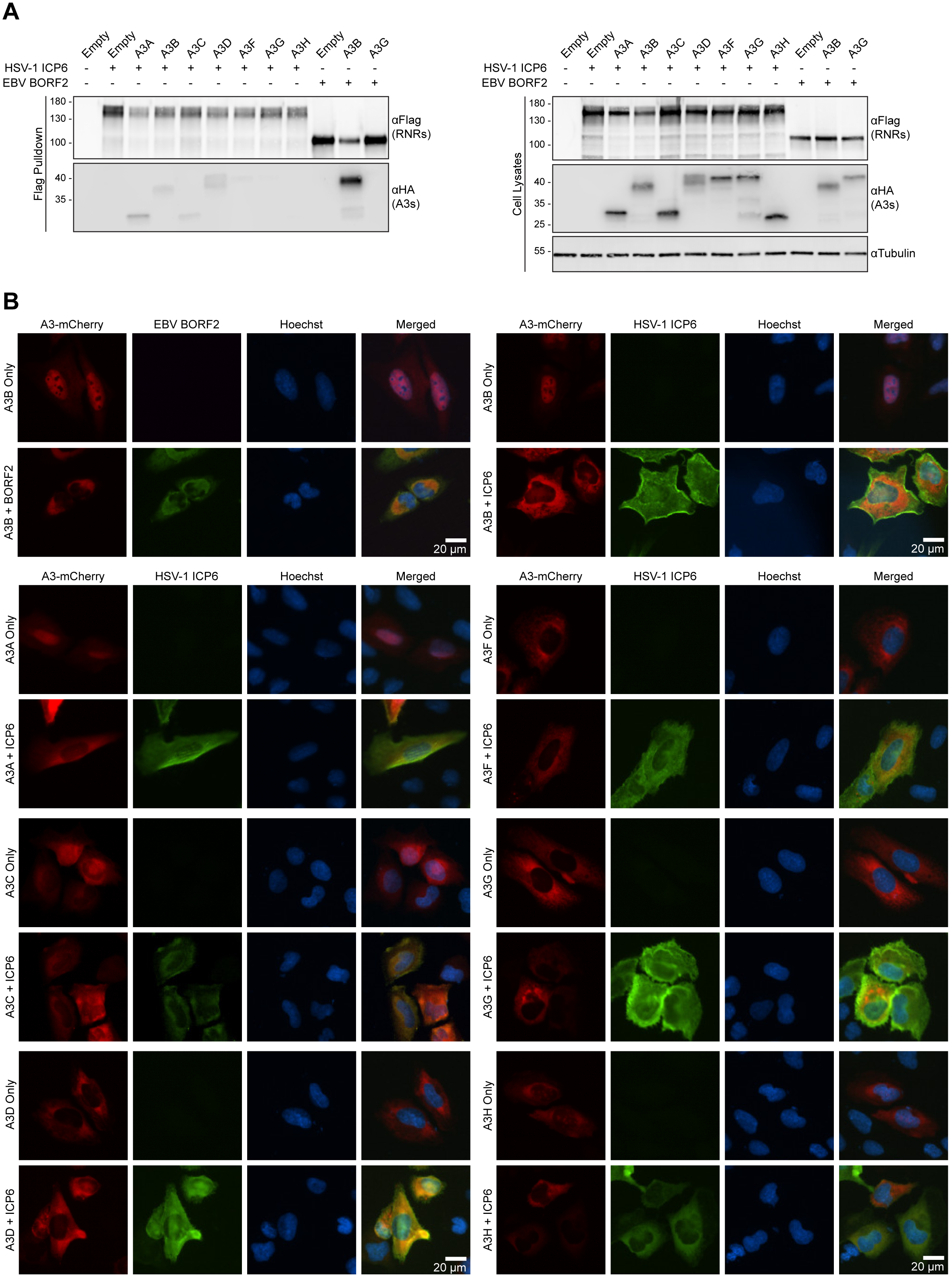
HSV-1 ICP6 binds and relocalizes A3B and A3A. (**A**) Co-immunoprecipitation of transfected HSV-1 FLAG-ICP6 with the indicated A3-HA constructs in 293T cells. Cells were lysed 48 hours post-transfection for anti-FLAG pulldown and resulting proteins were analyzed by immunoblot. EBV FLAG-BORF2 transfected with A3B and A3G were used as positive and negative co-IP controls, respectively. (**B**) Representative images of U2OS cells transfected with either A3-mCherry or FLAG-RNR constructs. Cells were fixed 48 hours post-transfection, permeabilized, and stained with anti-FLAG antibody and Hoechst. Co-transfection with A3B-mCherry and EBV FLAG-BORF2 was used as positive controls for relocalization from nuclear to cytoplasmic aggregates. A3 localization was compared in the presence and absence of HSV-1 FLAG-ICP6 co-transfection.

Next, IF microscopy was used to assess functional interactions between HSV-1 ICP6 and each of the human A3 enzymes. Human U2OS osteosarcoma cells were co-transfected with mCherry-tagged A3s and either empty vector or FLAG-tagged HSV-1 ICP6 and analyzed by IF after 48 hours (**Fig 4B**). On its own EBV BORF2 shows a cytoplasmic distribution and, as shown above and previously (18), it was able to completely relocalize A3B from the nucleus to cytoplasm. In comparison, HSV-1 FLAG-ICP6 showed a broadly cytoplasmic localization that did not change significantly with co-expression of any A3. However, co-expression of FLAG-ICP6 and A3B-mCherry or A3A-mCherry led to a near complete relocalization of these DNA deaminases from the nucleus to the cytoplasm. HSV-1 ICP6 did not cause significant relocalization of any of the other A3s. The dramatic relocalization results with A3B and A3A suggested that functionally relevant interactions may be occurring with these enzymes.

### HSV-1 infection relocalizes A3B, A3A, and A3C

To address whether HSV-1 infection similarly promotes relocalization of A3B and A3A, U2OS cells were transfected with A3-mCherry constructs 48 hours prior to either mock or HSV-1 infection. We used K26GFP, a HSV-1 strain that has a GFP moiety fused to capsid protein VP26 to allow for identification of infected cells (27). Cells were analyzed by IF microscopy 8 hours post-infection (hpi) (**Fig 5**). Similar to the ICP6 overexpression experiments described above, HSV-1 infection caused A3A to relocalize to the cytoplasmic compartment and A3B to change from a predominantly nuclear localization to a more cell-wide distribution. A3C also changed from a predominantly cytoplasmic localization to a more diffuse whole cell distribution, whereas A3D, A3F, A3G, and A3H were unchanged by HSV-1 infection. In an independent experiment, HSV-1-induced relocalization of A3A-mCherry and A3B-mCherry was quantified and compared to A3G-mCherry as a representative non-altered control (**Supplementary Fig 2**). This analysis confirmed that HSV-1 infection leads to significant changes in both A3A and A3B localization, whereas A3G is unaffected. Similar relocalization patterns were found in HeLa cells following HSV-1 K26GFP infection (**Supplementary Fig 3**). Moreover, time-course experiments showed that relocalization of A3A was detectable as early as 3 hpi, whereas A3B and A3C relocalization became apparent by 6 or 9 hpi (**Supplementary Fig 4**). These kinetic differences may reflect a differential affinity of the viral protein(s) to bind to these cellular A3 enzymes and/or different competitions with cellular interactors.

**Fig 5.**
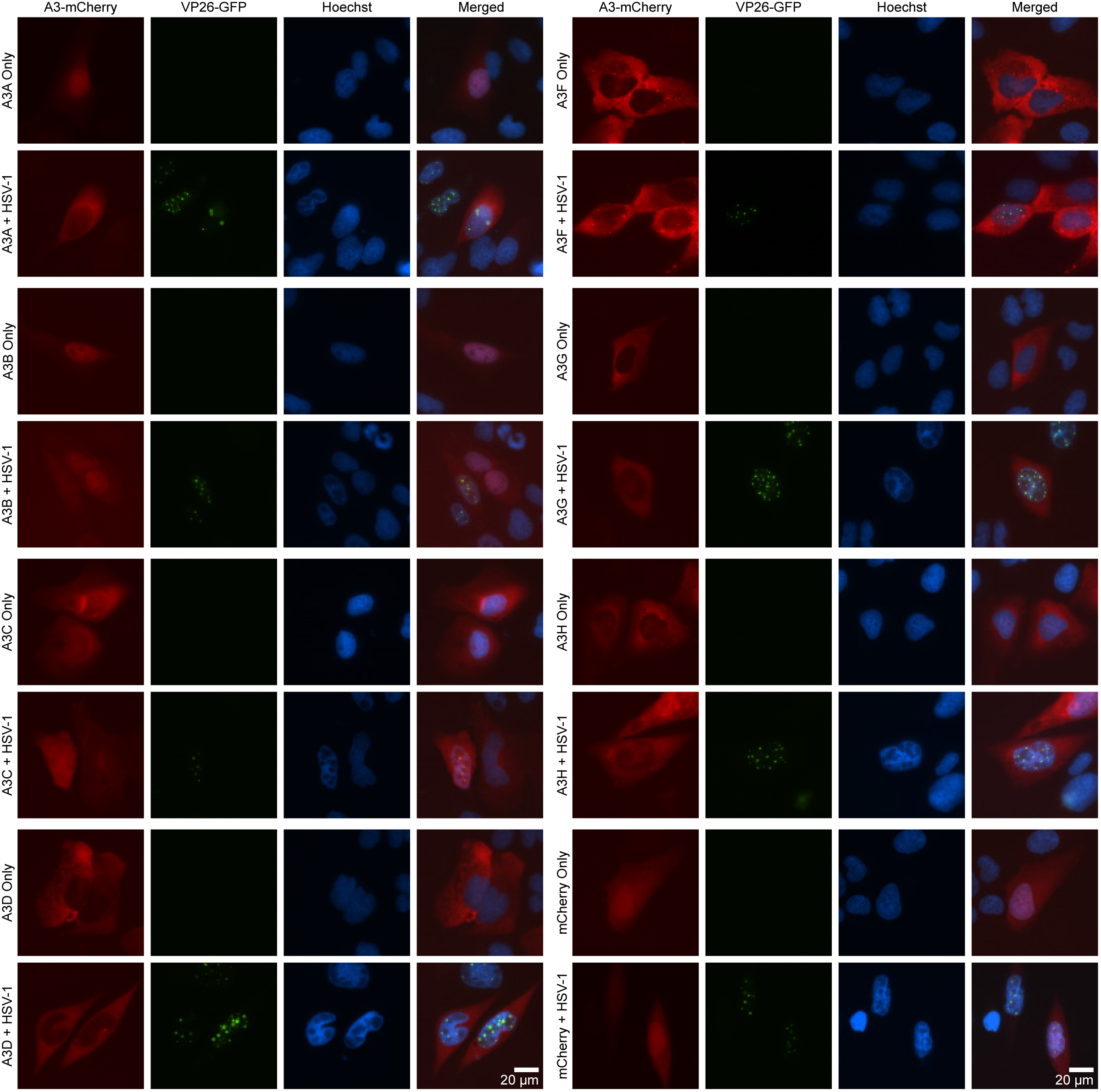
HSV-1 infection relocalizes A3B and A3A. Representative images of U2OS cells transfected with A3-mCherry constructs, followed by mock or HSV-1 K26GFP infection 48 hours post-transfection. Cells were fixed 8 hpi and stained with Hoechst, then imaged directly. The viral capsid protein VP26 is tagged with GFP which marks infected cells.

### HSV-1-mediated relocalization of A3B and A3A requires ICP6

To investigate whether the HSV-1 large RNR subunit is required for A3A/B/C relocalization, we next examined A3 localization in cells following infection with an HSV-1 KOS1.1 strain lacking ICP6 due to a deletion of the *UL39* gene (*UL39* encodes ICP6) (28). Vero cells were transfected with A3-mCherry constructs 48 hours prior to mock infection or infection with KOS1.1 or KOSΔICP6. After 8 hours, cells were fixed, permeabilized, and subjected to IF analysis by staining for the HSV-1 immediate early protein ICP27 to mark infected cells, and monitoring A3 localization through mCherry fluorescence. As above, HSV-1 infection caused the relocalization of A3A, A3B, and A3C (**Fig 6A**). However, only the relocalization A3A and A3B was ICP6-dependent, whereas A3C redistributed regardless of the presence of ICP6. Quantification of A3A and A3B relocalization showed that these proteins were not significantly changed upon KOS1.1ΔICP6 infection compared to mock-infected cells (**Supplementary Fig 2**). These results provide strong support for mechanistic conservation of the RNR large subunit interaction with A3A and A3B and also indicate that A3C relocalization by HSV-1 is mechanistically distinct.

**Fig 6.**
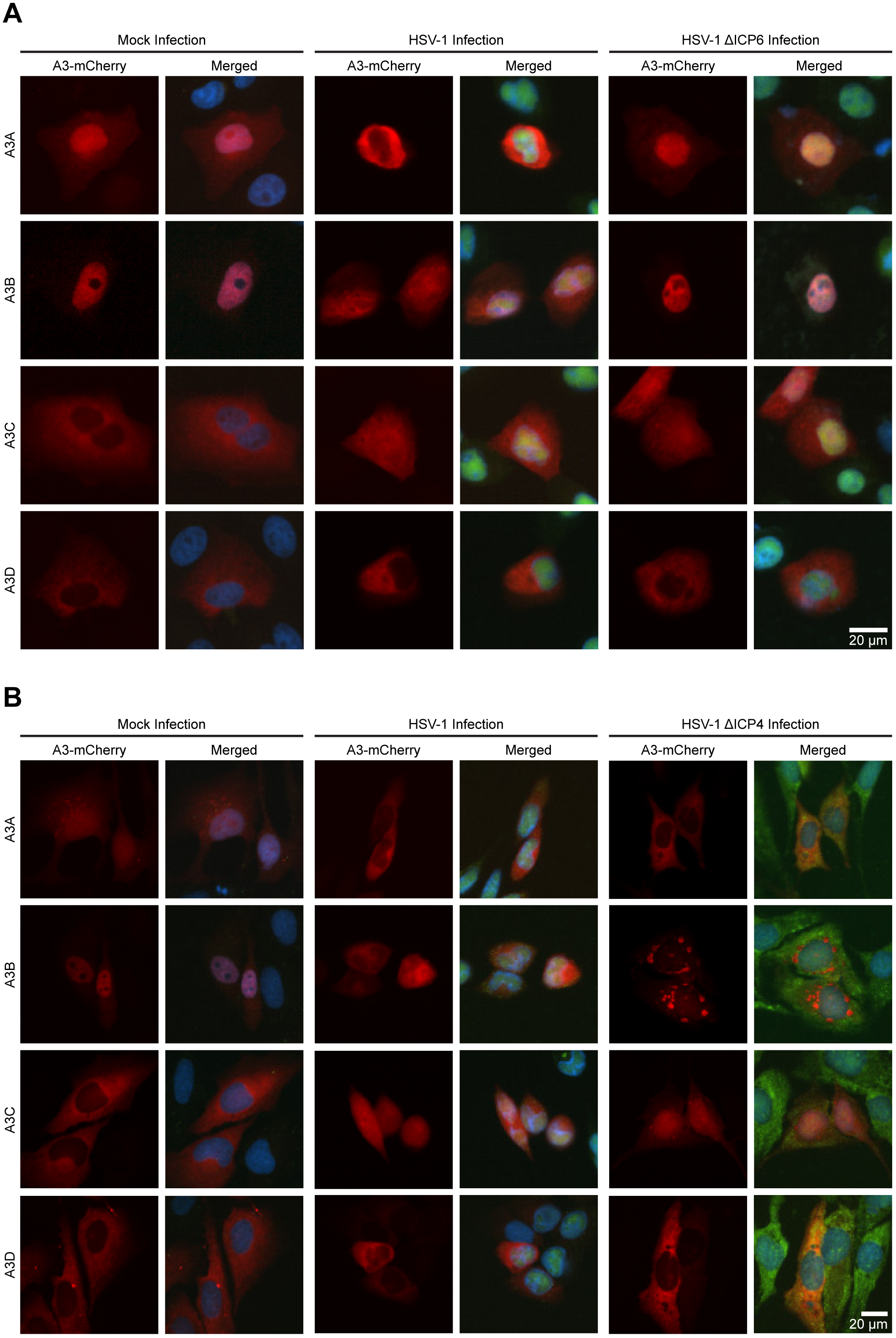
A3B and A3A relocalization is dependent on HSV-1 ICP6. **(A)** Representative images of Vero cells transfected with A3-mCherry constructs, followed by mock, wild-type HSV-1 KOS1.1, or HSV-1 KOS1.1ΔICP6 infection 48 hours post-transfection. Cells were fixed 8 hours after HSV-1 infection, permeabilized, and stained with anti-ICP27 antibody to mark infected cells and Hoechst. **(B)** Representative images from an experiment similar to that described in panel A, except using U2OS cells and the mutant virus HSV-1 KOS1.1ΔICP4.

To further investigate the role of ICP6 in mediating A3A and A3B relocalization, U2OS cells were infected with an HSV-1 KOS mutant with a deletion in the *ICP4* gene (29). ICP4, an immediate early protein, is the major transcriptional activator protein of HSV-1 (29). *ICP4*-null mutants exhibit a strict block to expression of nearly all viral delayed-early and late genes, but are competent to express the viral immediate-early genes (*ICP0, ICP22, UL54*, and *US12*) as well as the *UL39* gene, a delayed-early gene that is uniquely transactivated by ICP0 (30). In fact, at intermediate and late times post-infection, *ICP4*-null mutants express abnormally high levels of these immediate early proteins as well as ICP6 (29). Similar to what was seen for wild-type HSV-1 infection, infection with the HSV-1 KOSΔICP4 mutant also led to A3A and A3B relocalization, but with noticeably more pronounced phenotypes (**Fig 6B** and **Supplementary Fig 2**). For instance, this mutant virus caused A3B-mCherry to form perinuclear aggregates reminiscent of previously observed BORF2-A3B bodies (18) (**Fig 6B**). Interestingly, A3C localization became predominantly nuclear upon HSV-1 KOSΔICP4 infection, suggesting that one of the other four immediate early proteins besides ICP4 induces its relocalization. Taken together, these data show that HSV-1 ICP6 is both necessary and sufficient for the relocalization of A3A and A3B, and that at least one other viral factor is responsible for A3C relocalization. Identification of this factor will be the subject of a future investigation.

### Effect of A3B and A3A on HSV-1 replication

We next sought to test the effect of A3 expression on HSV-1 virus replication, with or without ICP6. HFF-1 cells were stably transduced to express HA-tagged A3 constructs and then infected at a low MOI (0.001 PFU/cell) with wild-type HSV-1 KOS1.1 or KOSΔICP6. At 48 hpi, the cultures were harvested, and after freeze-thawing to release infectious progeny, the cell lysates were titered on Vero cells to compare virus production. As previously described, KOSΔICP6 exhibited a 1-2 log defect in virus replication compared to wild-type KOS (28). However, there was no significant difference in either KOS1.1 or KOSΔICP6 virus titers produced from control HFF-1 cells or HFF-1 cells expressing different A3 family members (**Supplementary Fig 5A**).

To further test whether A3B or A3A can restrict HSV-1 replication, we performed plaque assays on either U2OS or Vero cells stably transduced with HA-tagged A3 constructs. Confluent monolayers were incubated with serial dilutions of KOS1.1 or KOSΔICP6 and incubated for 3 days to allow for plaque formation. However, expression of A3A or A3B did not have a discernable effect on the number or size of KOS1.1 or KOSΔICP6 plaques (**Supplementary Fig 5B**). These data suggest that even without ICP6, HSV-1 is not readily susceptible to restriction by A3A or A3B, possibly because it possesses other defenses against these virus restriction factors.

## Discussion

We previously described a novel mechanism for A3B counteraction by the γ-herpesvirus RNR large subunits, EBV BORF2 and KSHV ORF61 (18). These viral proteins interact directly with A3B, inhibit its DNA deaminase activity, and relocalize it from the nuclear to the cytoplasmic compartment. The importance of this A3B counteraction mechanism is evidenced by BORF2-null EBV eliciting lower viral titers, decreased infectivity, and an accumulation of A3B signature C/G-to-T/A mutations. Here, we investigated the question of specificity by comparing interactions with the full repertoire of seven different human A3 enzymes, and we also addressed the potential for broader conservation by asking whether the α-herpesvirus HSV-1 possesses a similar APOBEC3 relocalization mechanism. Although EBV BORF2 and KSHV ORF61 were able to interact with several different A3 proteins in co-IP experiments, these viral RNR large subunits only promoted the relocalization of A3B and A3A. HSV-1 ICP6 showed a similarly broad range of co-IP interactions but also only promoted the relocalization of A3B and A3A. Wild-type but not ICP6 deletion mutant HSV-1 infections yielded similar A3B and A3A relocalization phenotypes. These studies combine to indicate that human γ- and α-herpesviruses possess a conserved A3B/A relocalization mechanism mediated by the viral RNR large subunit.

The γ- and α-herpesvirus subfamilies encode both large and small RNR subunits (**Fig 1A**). These RNRs are thought to serve the canonical function of synthesizing deoxyribonucleotides by reducing the 2’-hydroxyl from ribonucleotide substrates (31). While RNRs are essential for all cellular life, the requirement for endogenous viral RNRs differs tremendously across viral families. For example, most small dsDNA viruses and single-stranded DNA viruses do not encode RNRs and instead rely on host-encoded RNRs for deoxyribonucleotide production (32, 33). On the other hand, RNRs are almost ubiquitous among large double-stranded DNA (dsDNA) viruses, such as herpesviruses and poxviruses, presumably due to high dNTP requirements during DNA replication (34-36). β-herpesviruses such as CMV are an exception, however, because they lack a small subunit and the large subunit has a defective catalytic site (37). In addition to ribonucleotide reductase activity, some viral RNRs have been shown to engage in non-catalytic activities that result in proviral phenotypes. For instance, the HSV-1 and HSV-2 large ribonucleotide reductase subunits, ICP6 and ICP10, respectively, have unique N-terminal extensions that block caspase-8 activity to inhibit apoptosis and bind RIP3 to promote necroptosis (38-41) (**Fig 1B**). CMV UL45 also has anti-apoptotic and pro-necroptotic functions suggesting this could be its predominant function (41-43).

The question of whether A3B, A3A, or both enzymes is most relevant to γ- and α-herpesvirus pathogenesis is likely to depend, at least in part, on the complex interplay between viral tropism(s) and alternating modes of latent versus lytic replication. For EBV, epithelial cells serve as the source of primary infection which are mandatory for establishing lytic replication cycles for person-to-person spread and enabling secondary infection of B lymphocytes for establishment of long-term latency (44). B cells also support lytic reactivation for reinfection and maintenance of EBV in the blood (45). Here, A3B may be more important than A3A simply because its expression is well-documented in these cell types (46, 47). Likewise, KSHV infects epithelial and B cells, but also engages in infection of clinically relevant endothelial cells which can lead to Kaposi’s sarcoma (48). Additionally, because monocytes are likely to be a secondary reservoir for KSHV infection (49-51), it is plausible that this virus requires the capacity to relocalize both A3B and A3A (A3B neutralization for replication in B cells and A3A neutralization for replication in monocytes/macrophages, where A3A is interferon-inducible and capable of being expressed at extremely high levels (46, 52, 53)). For HSV-1, although neither A3B nor A3A expression has been reported in neural/CNS cells, lytic replication in epithelial cells may require functional neutralization of A3B and/or A3A (54, 55).

The observation that the HSV-1 ΔICP6 mutant replicates at similar levels to wild-type HSV-1 in the presence of A3B or A3A was unexpected, but not entirely surprising. Given the large genomes of herpesviruses, it is possible that other viral proteins may have overlapping redundant functions in A3 counteraction and/or repair mechanisms to overcome A3-mediated hypermutation. One prime candidate is the viral-encoded uracil DNA glycosylase, encoded by the *UL2* gene, which has been shown to associate with the HSV-1 DNA polymerase in the infected cell nucleus (56). Consistent with this idea, we previously found that inhibition the EBV uracil DNA glycosylase (UDG) through expression of a universal UDG inhibitor (Ugi) results in enhanced A3B-mediated hypermutation of EBV genomes (18). It is thus possible that HSV-1 UL2 mediates the repair of uracil lesions generated by A3 enzymes allowing the virus to tolerate moderate levels of mutation in the absence of ICP6. It is also conceivable that HSV-1 encodes an additional, novel A3A/B neutralization or escape mechanism that is able to fully compensate for loss of ICP6 function (at least in the cell types tested here). Alternatively, inherent differences in viral DNA replication between HSV-1 and EBV could account for differences in replication phenotypes. HSV-1 replicates faster than EBV (57), which could result in less accessible single-stranded DNA for A3-mediated deamination. Lastly, the lack of an *in vitro* infectivity phenotype does not preclude *in vivo* disease relevance. Although prior studies have tested the impact of A3A and A3G (and APOBEC1) on wild-type HSV-1 replication in transgenic mice (58, 59), dedicated functional studies with mutants that at least partly cripple each viruses’ A3 relocalization mechanism(s) in the most disease relevant *in vivo* systems will be required to fully address the question of whether A3B, A3A, or both enzymes are relevant to the pathogenesis of these herpesviruses.

## Materials and Methods

### Generation of herpesvirus phylogenetic tree

Amino acid sequences for herpesvirus ribonucleotide reductase large subunits were obtained from NCBI Protein RefSeq with the following GenBank accession numbers: HSV-1 ICP6 YP_009137114.1, HSV-2 ICP10 YP_009137191.1, VZV ORF19 NP_040142.1, EBV BORF2 YP_401655.1, HCMV UL45 YP_081503.1, HHV6A U28 NP_042921.1, HHV6B U28 NP_050209.1, HHV7 U28 YP_073768.1, KSHV ORF61 YP_001129418.1. Alignment was generated using MUSCLE: multiple sequence alignment with high accuracy and high throughput (60) and phylogenetic tree was made using a neighbor-joining tree without distance corrections. Output was made using FigTree using scaled branches (61).

### DNA constructs for expression in human cell lines

The full set of pcDNA3.1(+) human APOBEC-HA expression constructs has been described (62) [A3A (GenBank accession NM_145699), A3B (NM_004900), A3C (NM_014508), A3D (NM_152426), A3F (NM_145298), A3G (NM021822), A3H (haplotype II; FJ376615)]. The full set of APOBEC-mCherry expression constructs was PCR amplified with Phusion High Fidelity DNA Polymerase (NEB M0530) from previously described A3-mCherry constructs (22) and subcloned into pcDNA5/TO (Invitrogen V103320). The forward PCR primers are as follows: A3A (5’-NNN NAA GCT TAC CAC CAT GGA AGC C-3’), A3B and A3C (5’-NNN NNA AGC TTA CCA CCA TGA ATC CA-3’), A3D (5’-NNN NNA AGC TTA CCA CCA TGA ATC CA-3’), A3F (5’-NNN NNA AGC TTA CCA CCA TGA AGC CT-3’), A3G (5’-NNN NAA GCT TAC CAC CAT GAA GCC T-3’), and A3H (5’-NNN NAA GCT TAC CAC CAT GGC TCT G-3’). The reverse PCR primer used was 5’-AGA GTC GCG GCC GCT TAC TTG TAC A-3’. PCR fragments were digested with *Hind*III-HF (NEB R3104) and *Not*I-HF (NEB R3189) and ligated into pcDNA5/TO. The full set of pLenti-iA3i-HA constructs were previously described except the puromycin resistance gene was replaced with a hygromycin resistance gene (63). Briefly, this is a lentiviral construct with an intron spanning the A3 gene with a C-terminal 3xHA tag, arranged in the antisense direction, which is expressed after reverse transcription and integration. This construct bypasses limitation of self-restriction by A3-mediated deamination of its own plasmid.

EBV BORF2 (GenBank accession V01555.2) with a C-terminal 3x-FLAG (DYKDDDDK) tag and EBV BaRF1 (Genbank accession V01555.2) with a C-terminal 3x-HA (YPYDVPDYA) tag was previously described (18). Other viral RNRs were subcloned with Phusion High Fidelity DNA Polymerase from previously described pCMV-3F vectors (18).

KSHV ORF61 (GenBank accession U75698.1) was PCR amplified using primers 5’-NNN NGA ATT CGC CAC CAT GTC TGT CCG GAC ATT TTG T-3’ and 5’-NNN NGA ATT CGC CAC CAT GTC TGT CCG GAC ATT TTG T-3’, digested with *Eco*RI-HF (NEB R3101S) and *Not*I-HF, and ligated into pcDNA4 with a C-terminal 3x-FLAG. The same construct was PCR amplified using primers 5’-NNN NGC GGC CGC GTC TGT CCG GAC ATT TTG T-3’ and 5’-NNN NTC TAG ATT ACT GAC AGA CCA GGC ACT C-3’, digested with *Not*I-HF and *Xba*I, and ligated into a similar pcDNA4 vector with N-terminal 3x-FLAG.

HSV-1 UL39 (GenBank accession JN555585.1) was PCR amplified using primers 5’-NNN NGA TAT CCG CCA CCA TGG CCA GCC GCC CAG CC-3’ and 5’-NNN NGC GGC CGC CCC AGC GCG CAG CT-3’, digested with *Eco*RV-HF (NEB R1395) and *Not*I-HF, and ligated into pcDNA4 (Invitrogen V102020) with a C-terminal 3x-FLAG (20). The same construct was PCR amplified using primers 5’-NNN NGC GGC CGC GGC CAG CCG CCC AGC CGC A-3’ and 5’-NNN NTC TAG ATT ACA GCG CGC AGC TCG TGC A-3’, digested with *Not*I-HF and *Xba*I (NEB R0145S), and ligated into a similar pcDNA4 vector with N-terminal 3x-FLAG.

### Human cell culture

Unless indicated, cell lines were derived from established lab collections. All cell cultures were supplemented with 10% heat-inactivated fetal bovine serum (Gibco 16140-063), 1x Pen-Strep (Thermo Fisher 15140122), and periodically tested for mycoplasma (Lonza MycoAlert PLUS LT07-710). No cell lines have ever been mycoplasma positive or previously treated. 293T and Vero cells were cultured in high glucose DMEM (Hyclone), U2OS cells were cultured in McCoy’s 5A media (Hyclone), and HeLa cells were cultured in RPMI 1640 (Corning).

### Co-immunoprecipitation experiments and immunoblots

Semi-confluent 293T cells were grown in 6-well plates and transfected with plasmids and 0.6 µL TransIT-LT1 (Mirus 2304) per 100 ng DNA in 100 µL serum-free Opti-MEM (Thermo Fisher 31985062). A titration series was performed to achieve roughly equivalent protein expression by immunoblot for the A3 panel and RNR homologue co-IP experiments. Growth medium was removed after 48 hrs and whole cells were harvested in 1 mL PBS-EDTA by pipetting. Cells were spun down, PBS-EDTA was removed, and cells were resuspended in 300 µL of ice-cold lysis buffer [150 mM NaCl, 50mM Tris-HCl, 10% glycerol, 1% IGEPAL (Sigma I8896), Roche cOmplete EDTA-free protease inhibitor cocktail tablet (Roche 5056489001), pH 7.4]. Cells were vortexed vigorously and left on ice for 30 minutes, then sonicated for 5 seconds in an ice water bath. 30 µL of whole cell lysate was aliquoted for immunoblot. Lysed cells were spun down at 13,000 rpm for 15 minutes to pellet debris and supernatant was added to clean tube with 25 µL resuspended anti-FLAG M2 Magnetic Beads (Sigma M8823) for overnight incubation at 4 °C with gentle rotation. Beads were then washed three times in 700 µL of ice-cold lysis buffer. Bound protein was eluted in 30 µL of elution butter [0.15 mg/mL 3xFLAG peptide (Sigma F4799) in 150 mM NaCl, 50 mM Tris-HCl, 10% glycerol, 0.05% Tergitol, pH 7.4]. Proteins were analyzed by immunoblot and antibodies used include mouse anti-FLAG 1:5000 (Sigma F1804), mouse anti-tubulin 1:10,000 (Sigma T5168), and rabbit anti-HA 1:3000 (Cell Signaling C29F4).

### HSV-1 infections and plaque assays

The HSV-1 strains used were wild-type strain KOS1.1 (64), K26GFP (27), ICP6 deletion mutant ICP6Δ (28), and the ICP4 deletion mutant d120 (29). Titers of viral stocks were determined by plaque assay on either Vero cells (KOS1.1, K26GFP, and ICP6Δ) or ICP4-complemented E5-Vero cells (65). HSV-1 infections were carried out as described (66). For microscopy experiments, cells were infected at a MOI of 5 PFU/cell. To assay HSV-1 replication in A3-transduced U20S cells, cells were infected at a MOI of 0.001 PFU/cell and incubated for 48 h, at which time a volume of sterilized milk equal to the volume of infected cell medium was added to each well, and the cells were frozen at −80°C. Infectious progeny virus was released by 3 cycles of freeze-thawing and titered on Vero cells. HSV-1 plaque assays were carried out in liquid media supplemented with 1% pooled normal human serum as previously described (66). For the HSV-1 plaque assays shown in **Supplementary Fig. 5**, U2OS or Vero cells were stably transduced with A3 constructs prior to carrying out plaque assays.

### IF microscopy

For IF imaging of transfected cells, approximately 5×10^4^ Vero, HeLa, or U2OS cells were plated on coverslips and after 24 hrs, transfected with 200 ng pcDNA4-RNR-3xFLAG, 200 ng pcDNA5/TO-A3-mCherry, or both. After 48 hrs, cells were fixed in 4% formaldehyde, permeabilized in 0.2% Triton X-100 in PBS for 10 minutes, washed three times for 5 minutes in PBS, and incubated in blocking buffer (0.0028 M KH_2_PO_4_, 0.0072 M K_2_HPO_4_, 5% goat serum (Gibco), 5% glycerol, 1% cold water fish gelatin (Sigma), 0.04% sodium azide, pH 7.2) for 1 hr. Cells were then incubated in blocking buffer with primary mouse anti-Flag 1:1000 overnight at 4 °C to detect FLAG-tagged RNRs. Cells were washed 3 times for 5 minutes with PBS, then incubated in secondary antibody goat anti-mouse AlexaFluor 488 1:1000 (Invitrogen A11001) diluted in blocking buffer for 2 hrs at room temperature in the dark. Cells were then counterstained with 1 µg/mL Hoechst 33342 for 10 minutes, rinsed twice for 5 minutes in PBS, and once in sterile water. Coverslips were mounted on pre-cleaned slides (Gold Seal Rite-On) using 20-30 µL of mounting media (dissolve 1g n-propyl gallate (Sigma) in 40 mL glycerol overnight, add 0.35 mL 0.1M KH_2_PO_4_, then pH to 8-8.5 with K_2_HPO_4_, Q.S. to 50mL with water). Slides were imaged on a Nikon Inverted Ti-E Deconvolution Microscope instrument and analyzed using NiS Elements.

For immunofluorescence imaging of HSV-1-infected cells, approximately 5×10^4^ Vero, HeLa, or U2OS cells were plated on coverslips and after 24 hrs, transfected with 200 ng pcDNA5/TO-A3-mCherry. After 48 hours, cells were infected with HSV-1 K26GFP, HSV-1 KOS1.1, HSV-1 KOS1.1ΔICP6, or HSV-1 KOS1.1ΔICP4 at MOI 5. Cells were fixed in 4% formaldehyde 8 hours post-infection and then IF studies proceeded as above. Time course experiments were fixed at either 3, 6, 9, or 12 hours post-infection. HSV-1 K26GFP experiments did not require primary or secondary antibody staining steps. Cells infected with HSV-1 KOS1.1 and mutants were incubated in primary antibody mouse anti-HSV-1 ICP27 H1113 (Santa Cruz sc69807) 1:1000 overnight at 4 °C to detected HSV-1-infected cells. Secondary antibody staining, counterstaining with Hoechst, mounting, and imaging proceeded as above.

### IF microscopy quantification

For quantification of A3 nuclear to cytoplasmic ratio, IF images were analyzed using Fiji software to obtain mean fluorescence intensities (MFI) of nuclear compartments determined by Hoechst stain outline and cytoplasmic compartments determined by cell outline. MFI values for each compartment were divided and plotted using Prism. Statistical analyses were performed using an unpaired Student’s t-test (n.s. = not significant with p>0.01).

## Acknowledgements

We thank Sandy Weller, Neal Deluca, and Prashant Desai for HSV-1 strains, M. Sanders and staff at the University of Minnesota Imaging Center for assistance with fluorescence microscopy, J. Becker for assistance with confocal microscopy, D. Ebrahimi for bioinformatics analyses of A3 expression in different cell types, and P. Southern for thoughtful comments and feedback on this manuscript. These studies were supported in part by NCI P01 CA234228 and funds from the University of Minnesota College of Biological Sciences and Academic Health Center (to RSH) as well as Canadian Institutes of Health Research (CIHR) project grant 153014 (to LF). NIH training grants provided salary support for AZC (F30 CA200432 and T32 GM008244) and MCJ (T32 CA009138). JY-M was supported by Secretaría Nacional de Educación Superior, Ciencia, Tecnología e Innovación (SENESCYT). GG is a scholar under the Horizon2020 program (H2020 MSCA-ITN-2015). V.D.O. is supported by Research Grants from the University of Turin (RILO18) and from the Italian Ministry of Education, University and Research – MIUR (PRIN 2015, 2015RMNSTA). RSH is the Margaret Harvey Schering Land Grant Chair for Cancer Research, a Distinguished McKnight University Professor, and an Investigator of the Howard Hughes Medical Institute. The funders had no role in study design, data collection and analysis, decision to publish, or preparation of the manuscript. RSH is a co-founder, shareholder, and consultant of ApoGen Biotechnologies Inc. The other authors have declared that no competing interests exist.

